# Genetically modified human type II collagen for N- and C-terminal covalent tagging

**DOI:** 10.1101/145136

**Authors:** Andrew Wieczorek, Clara K. Chan, Suzana Kovacic, Cindy Li, Thomas Dierks, Nancy R. Forde

## Abstract

Collagen is the predominant structural protein in vertebrates, where it contributes to connective tissues and the extracellular matrix; it is also widely used in biomaterials and tissue engineering. Dysfunction of this protein and its processing can lead to a wide variety of developmental disorders and connective tissue diseases. Recombinantly engineering the protein is challenging due to posttranslational modifications generally required for its stability and secretion from cells. Introducing end labels into the protein is problematic, because the N- and C-termini of the physiologically relevant tropocollagen lie internal to the initially flanking N- and C-propeptide sequences. Here, we introduce mutations into human type II procollagen in a manner that address these concerns, and purify the recombinant protein from a stably transfected HT1080 human fibrosarcoma cell line. Our approach introduces chemically addressable groups into the N- and Ctelopeptide termini of tropocollagen. Simultaneous overexpression of formylglycine generating enzyme (FGE) allows the endogenous production of an aldehyde tag in a defined, substituted sequence in the N-terminus of the mutated collagen, while the C-terminus of each chain presents a sulfhydryl group from an introduced cysteine. These modifications are designed to enable specific covalent end-labelling of collagen. We find that the doubly-mutated protein folds and is secreted from cells, while higher-order assembly into well-ordered collagen fibrils is demonstrated through transmission electron microscopy. Chemical tagging of thiols is successful, however background from endogenous aldehydes present in wildtype collagen has thus far obscured the desired specific N-terminal labelling. Strategies to overcome this challenge are proposed.

## Introduction

Collagen belongs to a large family of extracellular matrix proteins that provide mechanical and supportive functions in connective tissues. Given that collagen represents over 25% of protein in our bodies, disorders affecting the structural stability of collagen-containing tissues are wide-ranging, caused by more than 1000 mutations identified in 12 of the more than 28 different collagens that occur in vertebrates.^1^ Collagens are identified by their unique triple-helical structure, consisting of a repeating (Glycine-X-Y) motif, in which X and Y can be any amino acid but are frequently proline and hydroxyproline. The major fibrillar collagens are collagens type I, II and III, each of which forms fibrils that assemble into networks or fibres depending on tissue type. Of relevance to this work, type II collagen is the major structural protein of cartilaginous tissues, as well as being found in the vitreous of the eye, the inner ear, and intervertebral disks. Its enhanced digestion by native enzymes is associated with aging and is particularly severe in osteo- and rheumatoid arthritis.^2^, ^3^ In many tissues, collagen fulfils an important mechanical role, for example acting as the stress-bearing portion of articular cartilage.

Mechanical manipulation and characterization techniques can offer great insight into the function, and dysfunction, of collagen.^4-9^ While such studies have traditionally focussed on high-level collagen structures, in recent years there has been increasing focus on understanding collagen’s mechanics at the molecular level.^10-17^ Collagen molecules can be subjected to strain, and their resultant response characterized, by stretching these individual proteins by their ends. To do so in a controlled manner requires the ability to functionalize and capture the two ends of the protein in an orthogonal fashion.

Early labelling experiments addressed the unique globular termini in the precursor form of the protein, procollagen (Figure 1). Propeptides in types I and II collagen (removed on processing to the mature form of collagen) contain cysteines that enable chemical modification via their sulfhydryl groups.^10^, ^11^, ^15^ ^17^ The globular nature of these propeptides, however, can add mechanical lability to the protein, meaning that it may be difficult to distinguish signatures of force-induced changes in collagen’s triple helix from deformation of the propeptide domains. More recent experiments have utilized commercially available collagens, and have targeted the telopeptide ends of the protein for manipulation via telopeptide-specific antibodies.^13^ This approach may have low efficacy due to the partial removal of telopeptides in the pepsin digestion used to produce the collagen. Additionally, antibody linkages are mechanically weaker than other linking strategies, such as biotin-streptavidin or covalent interactions. The latter have been used to address one end of type I collagen or the two distinct ends of type III collagen, but in both cases these have used commercially available protein, which is not amenable to sequence manipulation for downstream structure-function studies.^13^, ^16^ There is a need for specific functionalization of collagen produced from a recombinant expression system, which would facilitate studies of how mutations disrupt the triple helix. This ability to chemically address collagen would enable detailed single-molecule mechanical studies, ideally to high forces, and also be applicable to fluorescence imaging via coupling to fluorophores or other labels of interest.

**Figure 1.**
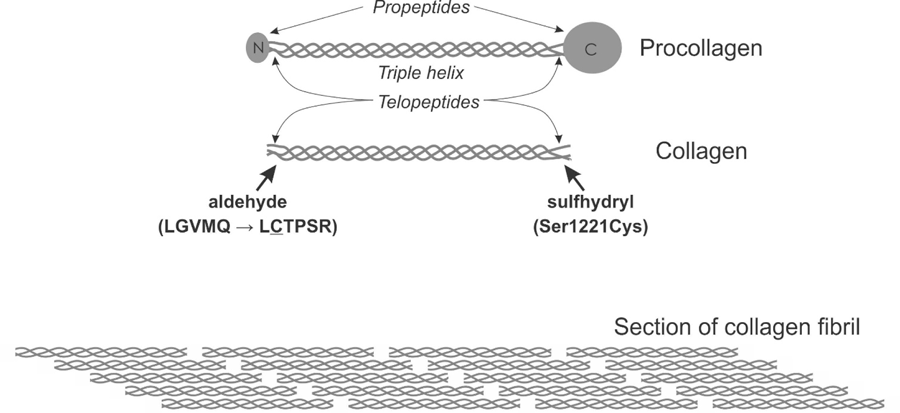
Schematic of collagen structure, with identification of the distinct forms (procollagen; collagen – also called tropocollagen; fibril) used in this work. The mutation strategy is illustrated, which aims to introduce orthogonal chemical handles into the telopeptides (terminus regions of collagen). The mutations were designed to minimize possible disruption to the triple helical structure while allowing the introduction of chemical tagging sites. In the N-telopeptide, our modified sequence substitutes the short sequence LGVMQ by LCTPSR, a sequence which is recognized by the enzyme formylglycine generating enzyme (FGE). In the C-telopeptide, we have introduced a Ser1221Cys mutation.

In this work, we describe the engineering of human type II procollagen to introduce chemically addressable moieties within the telopeptides (ends) of the mature form of the protein. Specifically, we describe the choice of mutations designed to introduce aldehyde and thiol groups to the two ends of collagen. We demonstrate that the mutated collagen is successfully secreted from the host human cell line, implying proper folding of the protein, and that it undergoes assembly into highly ordered fibrillar structures. Our experiments demonstrate challenges associated with the choice of an aldehyde functionality when using this cell line for procollagen modification, yet show successful chemical modification of the introduced thiols. This type II procollagen construct holds promise for future applications requiring tagging of the protein, including imaging and mechanical studies.

## Materials and Methods

*Mutagenesis:* Human type II procollagen constructs were produced with mutations in the N- and C-terminal telopeptides. Mutations were introduced at the plasmid level (Finnzymes Phusion site-directed mutagenesis kit, New England Biolabs) into the COL2A1-containing modified pYIC vector previously developed for stable expression of wild-type human procollagen.^17^ The plasmid contains an internal ribosome entry site (IRES) between open reading frames for COL2A1 and enhanced cyan fluorescent protein (ECFP), giving rise to uncoupled translation of both procollagen and this downstream fluorescent marker from a single mRNA transcript. This enables facile fluorescence-based screening of stably transfected mammalian cells. The Ntelopeptide sequence is modified by a substitution of the amino acid sequence LGVMQ by LCTPSR, a sequence which can be recognized by the enzyme formylglycine generating enzyme (FGE) to produce an aldehyde tag at the underlined cysteine.^18^ Primers used were 5’-CTG TGC ACC CCC AGC CGG GGA CCA ATG GGC CCC ATG GGA CCT CGA GG-3’ and 5’-CTG GGC GCC ACC AGC CTT TTC ATC AAA TCC TCC-3’. In the C-telopeptide, we introduced a Ser1221Cys mutation. Primers used were 5’-CCT GGC ATC GAC ATG TGC GCC TTT GCT GG-3’ and 5’-GCC AGG GGG ACC TGG AGG ACC AGG GG-3’. The resultant plasmid was linearized with Afl II and Rsr II to remove the neomycin resistance gene prior to transfection. *Stable cell line construction:* Unless otherwise mentioned, cloning, transformation, selection of stable cell lines and cell growth were performed essentially as described previously.^17^ In this work, an HT1080 fibrosarcoma cell line stably overexpressing the enzyme formylglycine generating enzyme (FGE)^19^, ^20^ was utilized for stable expression of the mutated procollagen. Endogenous expression of collagen IV by this cell line provides the necessary enzymes for correct posttranslational modification of the recombinant procollagen II.^17^, ^21^ To ensure stable retention of both introduced constructs (FGE and collagen), the HT1080 FGE-overexpressing cells (originally selected by G418 resistance)^19^, ^20^ were co-transfected with a linear puromycin resistance marker (Clontech 631626) and a linearized version of the plasmid DNA from which the neomycin selection was removed. After transfection, cells were selected in the presence of 400 μg/ml G418 and 0.25 μg/ml puromycin; surviving cells were screened for blue fluorescence indicating the expression of procollagen. Blue cells were tested for the secretion of procollagen. The introduction of mutations was confirmed by sequencing the RNA transcript from the stable cell line.

*Procollagen II purification:* Unless otherwise noted, purification of secreted procollagen II from media was performed as described previously.^17^ This process exploits two ion-exchange purification steps. The first step utilizes anion exchange with a DEAE column. The second purification step involves either further anion exchange (Q-sepharose) or cation exchange (Ssepharose). For cation exchange, the highest-concentration fractions of eluate from the initial DEAE column were dialysed into acetate buffer, pH 5.2 prior to loading on the column. Elution from the cation column was accomplished using a NaCl gradient, and detection and characterization of the collagen were performed as previously described. Purified protein was stored in 1X storage buffer (10 mM Tris HCl, 40 mM NaCl, 2.5 mM EDTA, pH 7.8) at 4°C. *Gel electrophoresis and Western blotting:* Protein samples were run on denaturing 6% polyacrylamide gels and were visualized by staining with Coomassie Blue. Western blotting to confirm identity used either a monoclonal antibody specific to type II collagen (Abcam Mab5B2.5) or an antibody cocktail against the triple helix region (Chondrex 7006).

*Fibril formation:* Fibril assembly proceeded essentially as described for wild-type collagen.^17^ Purified procollagen was dialyzed into phosphate buffered saline (PBS) pH 7.3 using Slide-ALyzer MINI Dialysis Units (20 kDa MWCO, Pierce) to a final estimated concentration of 150 μg/ml. To initiate fibril formation, 50 μl of this procollagen was incubated with 14 μl of 10 μg/ml Lys-C (Roche) at 37°C to remove N-and C-terminal propeptides.^22^ Collagen fibrils were isolated by centrifugation at 13,800g for 45 min. The supernatant was removed and the pellet gently resuspended in 10 μl of PBS.

*Transmission electron microscopy:* Samples were prepared by floating carbon-Formvar nickel grids (Ted Pella) on 5 μl of resuspended collagen fibrils in PBS for 1 hour. The grids were washed three times each with 50 μl of deionized water, blotted with Whatman filter paper and then negatively stained with 20 μl of 3% uranyl acetate (Ted Pella) for 45 seconds. Excess stain was removed by blotting and the grids were allowed to air dry at room temperature. The negatively stained samples were imaged at 200 kV accelerating voltage with a Hitachi 8000 transmission electron microscope at Simon Fraser University’s NanoImaging facility in 4D Labs. *Commercially obtained collagen:* Comparison experiments utilized acid-soluble type I collagen from rat tail tendon (Cultrex Invitrogen) and human type III collagen recombinantly expressed in yeast (Fibrogen).

*Chemical labelling:* Propeptides were cleaved from procollagen by incubation with 50 μg/ml chymotrypsin (Sigma, C7762) for four hours at room temperature.^17^ Following overnight dialysis at 4°C, collagen was purified using a Q-sepharose fast-flow column. Aldehyde-targeting reactions were performed with 5 mM hydrazide-biotin (Pierce EZ-link #21339) overnight at room temperature in acetate buffer (200 mM acetate buffer with 200 mM NaCl, pH 5.2). To target thiols, collagen was first reduced with 15 mM dithiothreitol (DTT) for 30 minutes at room temperature. Following dialysis into a reaction buffer (200 mM phosphate with 250 mM NaCl, pH 7.0), collagen was reacted with 0.3 mM maleimide-biotin (EZ-Link Maleimide-PEG2-Biotin, Thermo Scientific) at room temperature for one hour. Following each reaction, labelled collagen was dialysed into 1X storage buffer pH 8.0 and kept at 4°C. The success of either reaction was assessed by Western blotting for biotin with streptavidin-horseradish peroxidase.

## Results and Discussion

The aim of this work is to develop a system of production of human collagen that can be chemically addressed at its ends for future manipulation and imaging applications. Utilizing a recombinant form of the protein is advantageous, as it permits future investigations of the effects of point mutations on the structure and mechanics of the triple helix. To this end, we describe the modification of a previously developed recombinant expression system for type II human procollagen^17^ that introduces site-specific chemical labels within the so-called telopeptide termini of the mature form of the protein.

### Mutational strategy

Sequence screening was used to identify potentially permissive sites for the mutations described in the following paragraphs, which permit the use of covalent chemistry at each terminus. Figure 1 shows schematically the sites of the desired mutations. Both N- and C-terminal mutations are introduced at locations internal to Lys-C cleavage sites, such that the tags will remain following digestion of the propeptides by this endopeptidase^22^ or by chymotrypsin^17^ (the triple helix is resistant to digestion by most proteases). The tags are external to the triple helical domain to minimize potential impact on collagen folding, which can be of particular concern near the C-terminus.^23-25^

The residue most commonly used for site-specific targeting of proteins is cysteine. Cysteines offer easy access to chemical modification of proteins due to their sulfhydryl moieties. While cysteines are present in the propeptide domains of collagens and have been used for specific labelling of procollagen,^15^ the proteolytic removal of N- and C-terminal propeptides during processing into the mature form of collagen, tropocollagen, means that it is necessary to find or introduce targetable side-chains internally into the tropocollagen itself.

Type III collagen is unique among fibrillar collagens in that it possesses cysteines within its C-terminal telopeptide region, a feature that has been used previously for site-specific labelling of this protein.^16^ Type II collagen does not possess cysteines, requiring its mutation to introduce this functionality. We engineered a Ser1221Cys mutation in the C-terminal telopeptide region of type II collagen, with the aim of minimally disrupting the local structure and folding of the triple helix, while presenting the desired thiol chemical functionality to the terminus region of the mature protein.

For applications such as single-molecule force spectroscopy, distinct labelling of collagen’s two termini is desired. This distinction requires the use of two orthogonal chemical labels. To this end, we sought an additional chemical group that could be addressable in conditions maintaining properly folded protein structure, and which was compatible and economical in the context of the utilized human cell expression system. Aldehydes present such an advantage, offering ease of reactivity with hydrazide and hydrazide-labelled moieties in aqueous solutions and ambient temperatures.^26^ Aldehydes have previously been introduced into the N-terminus of collagens via reaction with the small molecule pyridoxal 5′-phosphate (PLP).^13^, ^16^ We propose an alternative strategy, which produces aldehydes endogenously. This would facilitate tagging in cell-based experiments.

Formylglycine generating enzyme (FGE) generates the catalytic formylglycine (FGly) residue in the active site of sulfatase enzymes by post-translational modification of a conserved cysteine within a short linear recognition sequence consisting of the minimal motif CxPxR.^18, 19, 27^ Pro- and eukaryotic FGEs have been used to introduce aldehyde functionality into recombinant proteins via FGly-modification of the cysteine within the target sequence inserted into various positions of the engineered protein. This has been done successfully in bacterial and mammalian expression systems.^27-30^ Eukaryotic FGE is localized in the endoplasmic reticulum,^20^ where initial posttranslational modifications of procollagen also occur.^31^ The co-localization of these proteins suggests the opportunity for FGE reactivity with a target sequence on procollagen. To enable this modification of collagen, we introduced the target sequence of LCTPSR, a sequence recognized by FGE,^18^ using it to replace the wild-type sequence of LGVMQ within the N-terminal telopeptide. The underlined cysteine is the residue targeted for enzymatic conversion to formylglycine.

### Cloning, expression and purification of mutated human procollagen

We cloned constructs and created stable human fibrosarcoma (HT1080) cells expressing type II procollagen, mutated at the N-terminus to present an FGE recognition sequence for aldehyde modification and at the C-terminus to introduce cysteines. The protein that should result is depicted schematically in Figure 1. To produce the type II procollagen, the mutated collagen was cloned into an HT1080 cell line engineered to stably overexpress FGE.^19^, ^20^ Stable cells were produced and characterized essentially as described previously for the wild-type protein in a standard HT1080 cell line.^17^

Type II procollagen was purified from the media, following the procedure described for the wildtype procollagen.^17^ This validated that the introduced N- and C-terminal modifications are tolerated and do not prevent proper folding and secretion.^31^

An additional test compared the efficacy of the conventional anion exchange to cation exchange for purification. Cation exchange purification, utilizing an acidic buffer, facilitates subsequent chemical labelling of the aldehyde moiety (see below), as it removes the need for an additional dialysis step. Although procollagen unsurprisingly eluted from the two different columns with different profiles, the yields from the two different approaches were similar.

For subsequent labelling reactions, purified procollagen was digested with chymotrypsin to remove propeptides.^17^ Figure 2 demonstrates the purity of the resulting protein.

**Figure 2.**
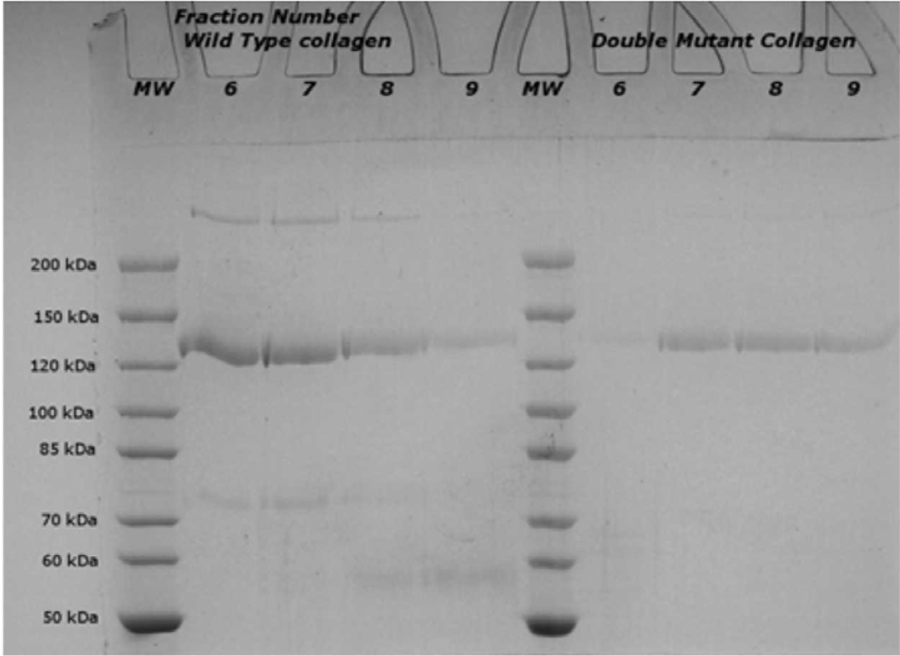
High purity of double-mutant type II collagen demonstrated by SDS-PAGE. The double-mutant collagen (obtained from a chymotrypsin digestion of the purified procollagen; right) runs as one band in a denaturing gel. The mobility is indistinguishable from that of the wildtype protein (left).

### Functional assessment: self-assembly into ordered fibrils

To verify function, we assayed the mutated collagen’s ability to form highly ordered collagen fibrils *in vitro*, which was assessed by imaging the fibrils with transmission electron microscopy (TEM). To generate a form of collagen capable of fibril formation, propeptides were removed by incubation with lysyl endopeptidase Lys-C, generating a form of collagen capable of self-assembly into well-ordered fibrils.^17^, ^22^ Fibril formation proceeded spontaneously in a solution of phosphate-buffered saline (PBS), pH 7.3, at 37°C.

TEM images of the resultant fibrils show that they display distinct light/dark D-periodic banding patterns (Figure 3). This is a distinguishing feature of well-ordered collagen fibrils. The periodicity is in the range of 65-70 nm, as expected.^32^, ^33^ These images demonstrate that the introduced mutations do not affect collagen’s ability to self-assemble into highly ordered fibrillar structures.

**Figure 3.**
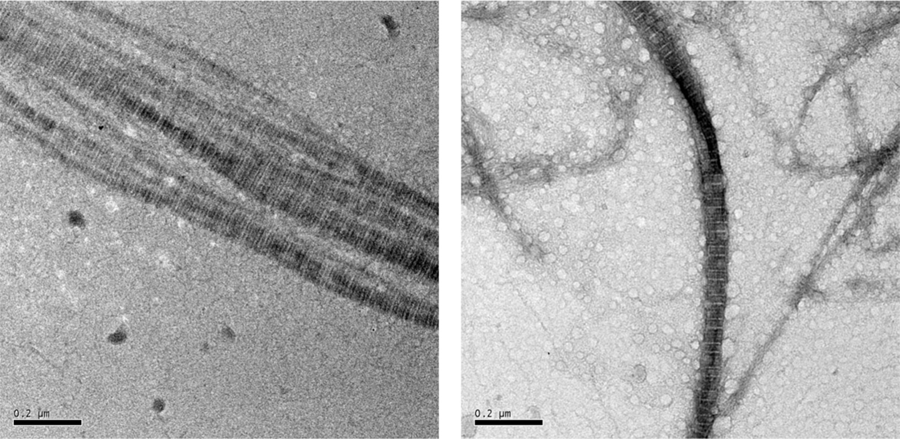
Transmission electron microscopy (TEM) imaging demonstrates that the double-mutant type II collagen can self-assemble to form well-ordered fibrils, as exemplified by the periodic banding pattern. Two representative images of the resulting fibrils are shown. Scale bars = 200 nm.

### Chemical labelling

The mutations introduced into the N- and C-telopeptides aimed to present orthogonal functional groups for covalent modification. While sequence analysis confirmed the presence of the mutations, it remained to be determined whether the N-terminal FGE sequence had been enzymatically modified to present an aldehyde, and furthermore, whether or not the functional groups were chemically accessible within the folded, secreted protein.

Accessibility of thiols from the cysteine residues was probed by reaction with biotin-maleimide. To eliminate the possibility of unwanted reactions with sulfhydryl groups of cysteines endogenously present in collagen’s propeptide regions, the propeptides were removed by chymotrypsin digestion. Following reduction of the resulting collagen to generate thiols (see Methods), we observed that the double-mutant collagen indeed reacted with biotin-maleimide, as it produced a strong signal in a Western blot with streptavidin (Figure 4). By contrast, similarly treated wildtype protein gave no signal, demonstrating that thiol functionality is a result of the introduced mutations. Because cysteines were introduced into each telopeptide end of collagen, the observed reactivity with maleimide does not assay for conversion of the C-terminal cysteine into formylglycine, but does demonstrate the successful introduction of reactive thiols.

**Figure 4.**
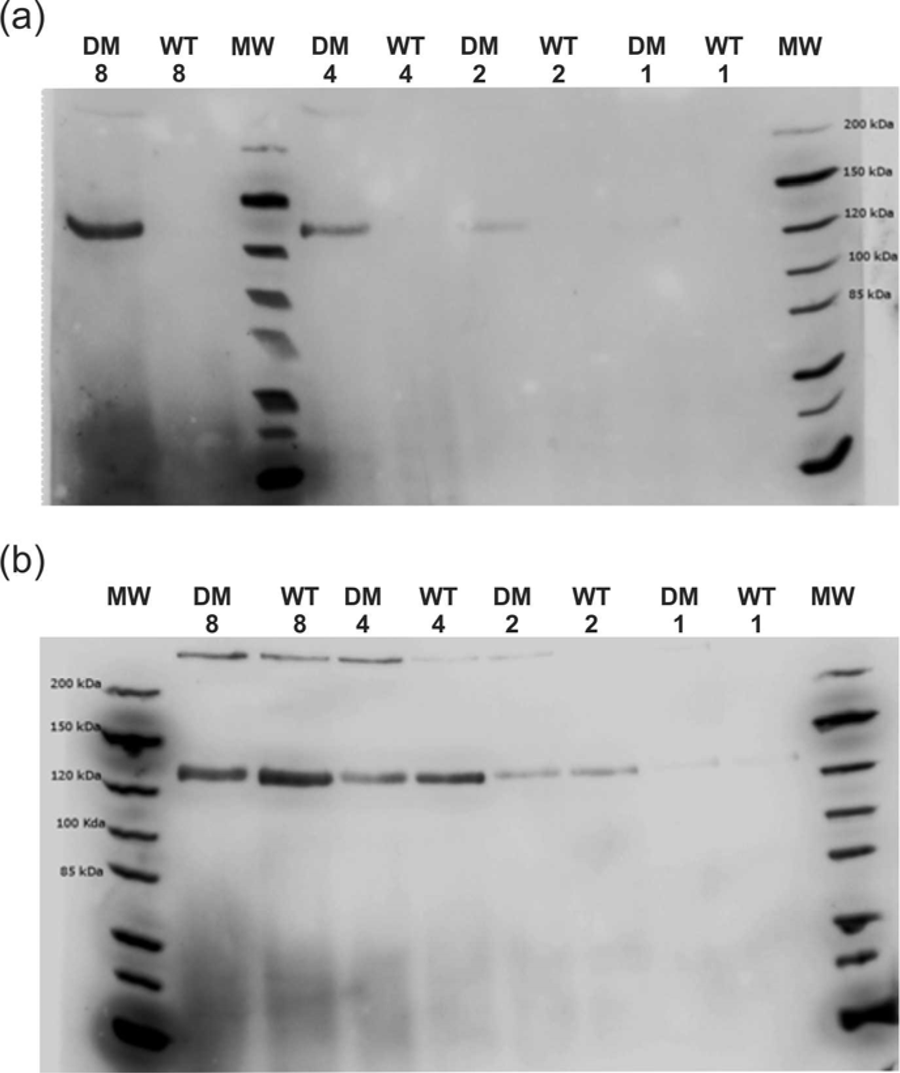
Western blotting shows that the double-mutant type II collagen is successfully labelled using maleimide-biotin. Following reaction of chymotrypsin-digested procollagen with maleimide-biotin, (a) the presence of biotin was probed for with streptavidin, while (b) blotting with an anti-collagen antibody cocktail confirmed similar amounts of double-mutant (DM) and wild-type (WT) collagen were loaded. Numbers indicate the relative amounts loaded in each lane. Biotinylation of sulfhydryls is specific to the double-mutant collagen.

To assay for presence and accessibility of the N-terminal aldehyde, we reacted the purified protein with biotin-hydrazide.^28^ Success of this reaction should demonstrate both that the cysteine within the FGE recognition sequence was enzymatically converted to aldehyde, and that it remains accessible to subsequent chemical labelling following folding and secretion. Unfortunately, these results demonstrated roughly equal reactivity of the double-mutant and wild-type proteins with hydrazide (Figure 5). Many replications of these results, under different reaction conditions, led us to conclude that aldehyde groups are present not only in the double-mutant but also in the wild-type recombinant protein. 12

**Figure 5.**
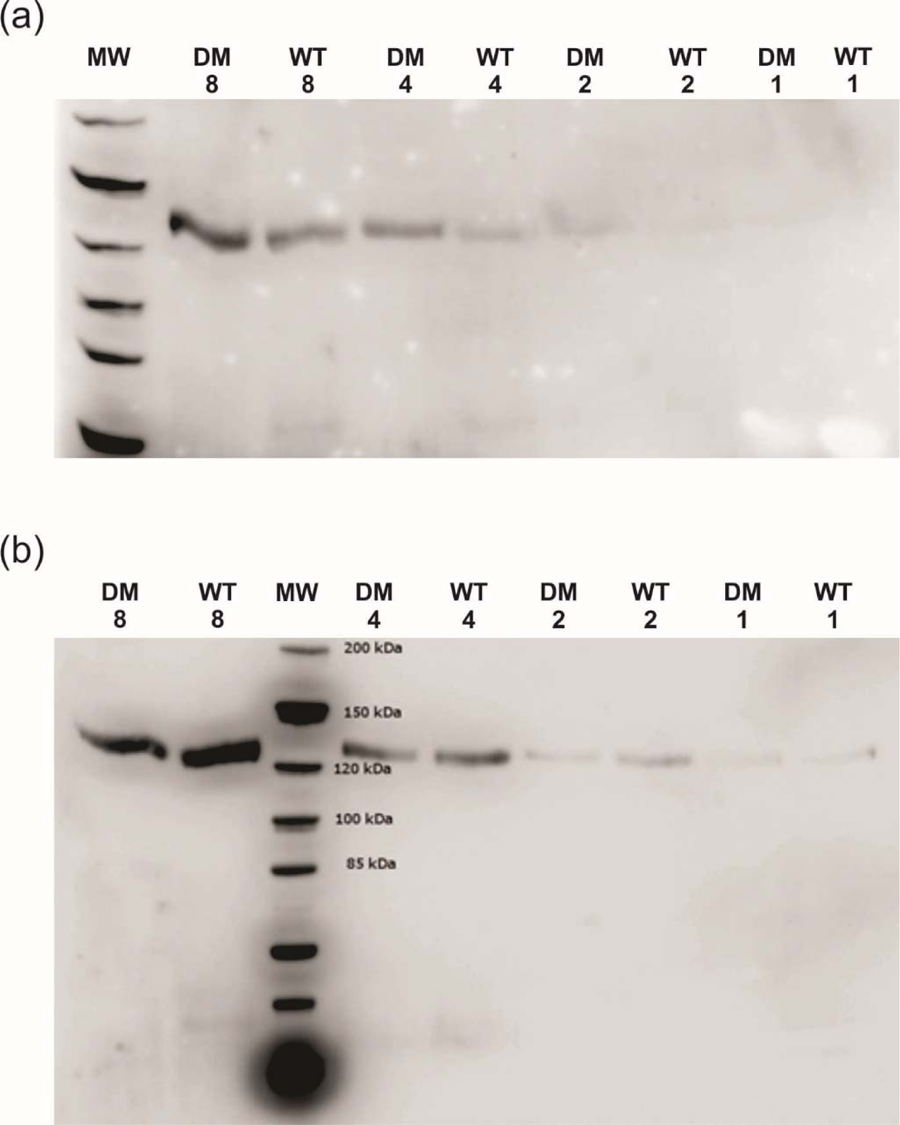
Western blotting shows that both the double-mutant and wild-type recombinantly expressed type II collagen are biotinylated, following reaction targeting aldehydes with hydrazide-biotin. (a) The presence of biotin was probed for with streptavidin, and (b) blotting with an anti-collagen antibody cocktail confirmed similar amounts of double-mutant (DM) and wild-type (WT) collagen were loaded. Numbers indicate the relative amounts loaded in each lane. Results indicate that aldehydes are present in both the double-mutant and wild-type collagen.

It appears that, contrary to our expectation, molecular collagen can undergo post-translational modification to introduce aldehydes. It is well known that as part of its production, assembly and secretion pathway, collagen undergoes extensive posttranslational processing.^34^ In the endoplasmic reticulum, prolines are hydroxylated and lysines hydroxylated, with hydroxylysines acting as subsequent targets for O-linked glycosylation.^34^ These modifications have been identified also in our wild-type recombinant procollagen.^17^ Aldehyde-producing posttranslational modifications in collagen have been attributed to lysyl oxidase enzymes, which act to convert lysine and hydroxylysine to allysine and hydroxyallysine, *i.e.*, their peptidyl aldehyde counterparts.^34^ This activity has generally been believed to occur extracellularly, stimulated by association of collagen into its fibrillar form.^34^ Aldehydes subsequently react to create crosslinks that stabilize fibrillar collagen.^35^ While the presence of aldehydes in tissue-derived collagen is not surprising, it was unexpected to discover them within our recombinantly expressed procollagen. So either allysine and/or hydroxyallysine were generated extracellularly, independent of stable fibril formation, or lysyl oxidase re-enters the cell and oxidizes collagen intracellularly.^36^ The study of collagen processing could benefit from the use of human-cell-based expression systems such as this,^37^ which are amenable to manipulation. Care must be taken, however, to infer too much from results utilizing a cell line that does not endogenously produce high levels of fibrillar collagens.

To provide additional evidence that the observed reactivity with hydrazide was due to aldehydes, we examined hydrazide reactivity of collagens from other sources that have different types of post-translational modifications. We reacted hydrazide-biotin with commercially available collagens, type I from rat tail tendon and human type III recombinantly expressed in yeast cells. Based on the discussion of the previous paragraph, we anticipated that the tissue-derived sample would possess aldehydes and that the collagen recombinantly expressed in yeast, which lacks most of the enzymes required for posttranslational processing of collagen,^38^ would not. Our results confirmed these expectations (Figure 6): type I collagen from rat reacted with hydrazide-biotin, while collagen produced from yeast did not. The yeast-derived collagen did, however, react with hydrazide-biotin following incubation with pyridoxal 5′-phosphate (PLP), which has been shown to modify N-termini of proteins to aldehydes.^39^ If yeast-expressed collagen, produced with only prolyl hydroxylation as a post-translational modification, is sufficient for experiments, then aldehyde modification via the PLP small-molecule reaction offers easy access to labelled collagen.^13^, ^16^ However, if collagen harboring a more complete suite of modifications is desired, a different cell line must be used.^38^ Based on the findings of this work, we propose that inhibition^34^ or a knockout of lysyl oxidase in a human-cell-based recombinant expression system could result in the ability to introduce aldehydes in collagen uniquely located within the FGE recognition sequence and available for specific chemical tagging. This remains to be tested.

**Figure 6.**
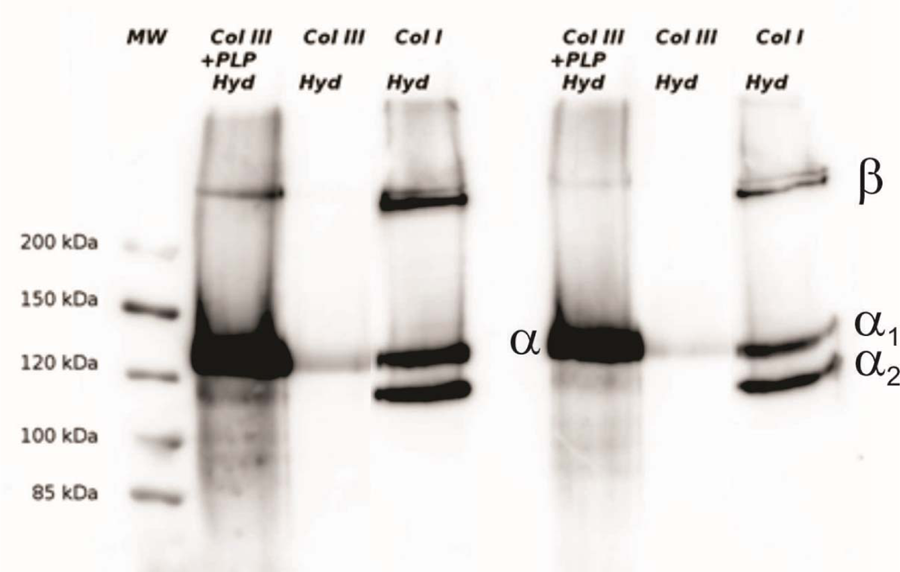
Western blotting with streptavidin-HRP demonstrates the differences in composition between tissue-derived and yeast-expressed collagen. Type I collagen from rat tail (Col I) reacts with biotin-hydrazide (Hyd), demonstrating the presence of endogenous aldehydes in this sample, while human type III collagen produced recombinantly in yeast (Col III) does not. Following incubation of this type III collagen with the small molecule PLP to introduce aldehydes (Col III + PLP), the protein becomes reactive to biotin-hydrazide. Replicates in the right lanes are at half the loading concentration as on the left. α_1_ and α_2_ refer to the two component chains of the heterotrimeric (α_1_)_2_(α_2_)_1_ type I collagen triple helix, while type III collagen, like type II, consists of three identical α chains. β bands refer to two crosslinked α-chains. These are a standard feature seen in SDS-PAGE gels of collagen and predominantly arise from crosslinking between chains within a triple helix.

## Conclusions

In this work, we have demonstrated that type II procollagen with specifically engineered mutations within its N- and C-telopeptide regions is capable of folding, secretion from cells, and self-assembly into well-ordered fibrils. The locations of these mutations were chosen to minimize sequence disruption, while providing unique chemically addressable groups at the end regions of the mature collagen protein. We have demonstrated that cysteines introduced through these mutations are accessible to modification by maleimide. Though the surprising presence of endogenous aldehydes in the recombinantly expressed proteins prevented the establishment of specificity and accessibility of the introduced aldehyde-producing mutation at the N-terminus of the protein, it did provide additional insight into collagen maturation in eukaryotic expression systems. Suggestions for future approaches to overcome this problem were provided.

## Acknowledgements

This research was funded by a Bone Health Catalyst Grant from the Canadian Institutes of Health Research (CIHR), by the Michael Smith Foundation for Health Research (MSFHR) and by a Discovery Grant from the Natural Sciences and Engineering Research Council of Canada (NSERC). We thank Veronica Tsai for assistance with sequencing.

